# The Role of Physical Activity in Mitigating Age-Related Changes in the Neuromuscular Control of Gait

**DOI:** 10.1101/2024.10.28.620570

**Authors:** M.N. Nuñez-Lisboa, A.H. Dewolf

**Affiliations:** Laboratory of biomechanics and Physiology of Locomotion, Institute of NeuroScience, Université catholique de Louvain, Place Pierre de Coubertin 1, Louvain-la-Neuve, 1348, Belgium; Exercise Science Laboratory, School of Kinesiology, Faculty of Medicine, Universidad Finis Terrae, Santiago, Chile

**Keywords:** Neuromuscular control, Gait mechanics, Physical activity, Aging

## Abstract

Exercise is known to induce several neural and muscular adaptations, such as increased muscle mass and functional capacity in older adults. In this study, we investigated its impact on the neuromuscular control of gait among young and older adults, divided into two groups: more active (young: n=15; 5185 ± 1471 MET-min/week; old: n=14; 6481 ± 4846 MET-min/week) and less active participants (young: n=14; 1265 ± 965 MET-min/week; old: n=14; 1473 ± 859 MET-min/week). Maximal isometric tests of ankle and knee extension revealed a reduction in force among older adults, with differences associated with the level of physical activity at the ankle level. Gait mechanics revealed no significant differences between young adults and the more active older adults. In contrast, less active older adults exhibited shorter steps, higher mechanical cost, and greater collision at heel strike. These changes cannot be attributed solely to reductions in muscle strength. Instead, they are likely the result of modifications in neuromuscular control and mechanical properties of muscles in less active older adults. Specifically, wider activation (and greater coactivation) of lumbar and sacral motor pools as well as a different timing of activation were observed. Also, their muscle-tendon stiffness was reduced. In conclusion, our findings highlight that the age-related decline in gait efficiency is exacerbated by a sedentary lifestyle. Even modest increases in physical activity appear to preserve neuromuscular control and improve walking performance. This suggests that interventions aiming to enhance physical activity levels could mitigate age-related declines in gait mechanics.

## Introduction

With extended life expectancy, the loss of mobility of older adults has dramatic individual and societal impacts. Among the many factors of frailty involved (e.g. metabolic, hormonal, immunological), loss of muscle mass and strength (Hyatt *et al*., 1990; Akima *et al*., 2001), changes in connective tissue properties (Graça *et al*., 2023), and neurodegenerative changes (Rygiel *et al*., 2016) play a key role. These neuromuscular alterations result in an increased metabolic rate of locomotion in older adults, associated with a loss of independence and an increased risk of morbidity and mortality (Guralnik *et al*., 1995; Cooper *et al*., 2010, 2011; Studenski *et al*., 2011).

Several age-related modifications of the muscle and tendon tissues have been described, such as loss of muscle fibres (Lexell *et al*., 1983; Lexell, 1995; McPhee *et al*., 2018), a shift in muscle fibre type distribution (Williamson *et al*., 2000), modification of the tendon and muscles cross-sectional area (Lexell & Taylor, 1991; Stenroth *et al*., 2012) or changes in the stiffness and viscoelastic properties of tendon and muscles (Stenroth *et al*., 2012; Akagi *et al*., 2015; Lim *et al*., 2019; Şendur *et al*., 2020; Çekok *et al*., 2024), associated with histological changes such as impaired connective tissues or myosteatosis. The decrease in muscle-tendon unit performance has an impact on mobility of the older adults. For example, older adults redistributed lower extremity joint moment and power compared with young people in walking (DeVita & Hortobagyi, 2000; Silder *et al*., 2008), so-called biomechanical plasticity. The scientific community (Winter *et al*., 1990; DeVita & Hortobagyi, 2000; Kerrigan *et al*., 2000; McGibbon, 2003; Franz & Kram, 2013; Franz, 2016; Gueugnon *et al*., 2019; Delabastita *et al*., 2021; Boyer *et al*., 2023) most often points to a reduction in mechanical power generated by the plantar flexor muscles during the push-off phase of walking as the hallmark biomechanical ageing features of gait.

The reduction of ankle muscle contractile performance has an impact on mobility of the older adults, but further limitations to the mobility come also from other factors (Marcucci & Reggiani, 2020). For instance, another determinant of functional capacity and autonomy is the integrity of other components of the neuromuscular system (Aagaard *et al*., 2010; Hunter *et al*., 2016). Indeed, we rely on the firing characteristics of our spinal motoneurons to produce force, and so, to perform daily activities like walking. Elderly adults have 30% fewer motor units innervating lower limb muscles than young adults (Tomlinson & Irving, 1977; McNeil *et al*., 2005), with denervation exceeding compensatory reinnervation by alpha motor neurons, muscle atrophy is further exacerbated (Hepple & Rice, 2016). Additionally, older adults exhibit lower peak firing frequencies during submaximal isometric contractions compared to younger adults (Hassan *et al*., 2021; Orssatto *et al*., 2022), contributing to a decline in muscle function.

Following the death of motoneurons, the nervous system displays astounding plasticity (Heppe *et al*., 2016; McNeil & Rice, 2018), with age-related changes in the organization of the motoneuron activity at the spinal cord level (Monaco *et al*., 2010; Dominici *et al*., 2011; Ivanenko *et al*., 2013; Dewolf *et al*., 2021*b*, 2021*c*). The age-related modifications of the spinal segmental output involve a widening of the patterns of activity of motoneurons, increased muscle co-activation, and a differential impairment of caudal versus rostral segments (Dewolf *et al*., 2021*b*, 2021*c*; Nùñez-Lisboa *et al*., 2023), which in turn can evoke alterations in muscle coordination and locomotor behaviour. Decoding the change in spinal motoneuron output in the elderly may be essential for linking the modification of output with neuromuscular degeneration and the firing characteristics of motoneurons.

High physical activity level is crucial for enhancing physical fitness by increasing muscle strength, and cardiopulmonary function and stimulating metabolic activity (Kraemer *et al*., 2002; da Cunha *et al*., 2015; Pour *et al*., 2017; Chen *et al*., 2021; el Hadouchi *et al*., 2022; Markov *et al*., 2022), decreasing risk of disability (Reid & Fielding, 2012). Practice and enhanced physical activity not only increase/maintain muscle mass, strength, power and functional capacity in older adults, but it also modify age-related changes in motor unit structure and function across the life span (Hunter *et al*., 2016; Cogliati *et al*., 2020). For example, exercise promotes various metabolic responses and morphological reconfigurations in the neuromuscular junction of older adults (Rogers & Nishimune, 2017), decelerating or even reversing neuromuscular junction degeneration (Wang *et al*., 2024). On the contrary, less regular physical activity among old adults may exacerbate age differences between young and older adults, for example in muscle activation (Harridge *et al*., 1999). However, enhancing physical capacity alone may not be sufficient to mitigate the age-related decline of the neuro- muscular system. Indeed, increasing the physical activity of older adults via resistance training fails to directly translate to improved propulsive power generation in walking (Beijersbergen *et al*., 2013). Similarly, Brach et al. (2013) assert that multicomponent impairment-based walking exercises, while beneficial for strength, flexibility, and endurance, don’t always lead to better walking ability. Also, a higher level of physical activity in older adults did not mitigate the age-related modification of kinematic coordination and distal-to-proximal redistribution (Boyer *et al*., 2012), suggesting that there remains a gap in understanding the biomechanical adaptations that physical activity levels may induce in gait mechanics across aging (Beijersbergen *et al*., 2013).

While a large body of literature has focused on the role of physical activity in preserving mobility, it is still difficult to disentangle the influence of biological aging from that of physical activity on the subsequent changes in the neuromuscular system. To address this question, we recorded data from both older and younger participants, divided according to their weekly physical activity levels. After evaluating the isometric strength and the mechanical properties of ankle and knee extensors, we then compared the mechanics and neuromuscular control of gait. The purpose of this study was to shed light on contributors to the age-related changes in biomechanical, physiological, and motor control adaptations of gait. We first hypothesized that a clear effect of aging on muscular mechanical properties, maximal isometric forces and neuromuscular control of gait would be observed. Also, we tested the hypothesis that more active older individuals exhibit a “slowing down” in the natural aging process of the neuromuscular system.

## Methods

### Participants

Fifty-nine healthy young and older adults (♂ = 39, ♀ = 20) participated in this study. Both age groups were divided into two groups (less and more active), according to the median level of physical activity, measured through the Global Physical Activity Questionnaire (GPAQ) (Fig. 1). A modified z-score using the median and Median Absolute Deviation (MAD) was applied to provide a robust classification of participants into “Less Active” and “More Active. The four groups’ characteristics are presented in Table 1. The number of subjects chosen was based on an a priori statistical power analysis using G_Power (version 3.1.9.7, Germany) (Faul *et al*., 2007) from the results of a previous study that focused on similar variables (Dewolf *et al*., 2021*c*), indicating that 12 subjects per group would be sufficient. The inclusion criteria were the following: ability to walk a kilometre, no locomotor system injury complaints and no previous history of neurological disorders. All participants provided informed consent, and the procedures used for this study were approved by the ethics committee of the Université Catholique de Louvain (Belgian Registration Number: B403201524765) and adhered to the Declaration of Helsinki.

**Figure 1.**
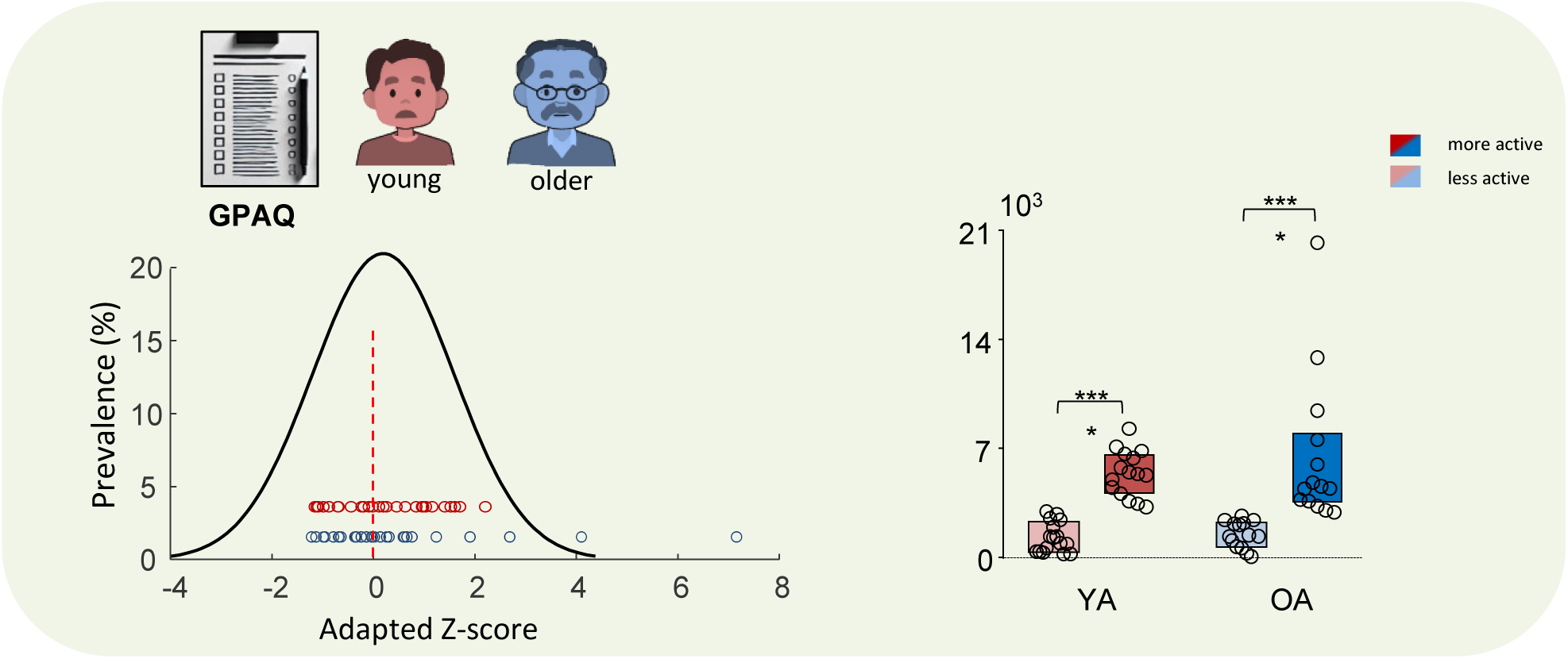
Group classification based on physical activity levels. *Left panel:* Distribution of physical activity levels in MET-min/week in young (red dots) and older adults (blue dots), classified using the Global Physical Activity Questionnaire (GPAQ). Older adults had a broader distribution of activity levels compared to the younger group. An adapted Z-score was applied to normalize activity levels. In this way, participants with a positive/negative adapted Z-score were classified as more/less active. *Right panel:* Boxplot displaying the average MET-min/week scores per group. The statistical significance of the differences between groups was assessed using the Kruskal-Wallis test. The asterisks indicate post hoc comparisons between groups (*p < 0.05, **p < 0.01, ***p < 0.001, ****p < 0.0001).

### Experimental procedure and data collection

The participants were first asked to stand on an instrumented treadmill to perform a static balance test. They stand with their feet together, closed, for 30 seconds each. During the test, they were instructed to maintain their arms relaxed at their sides, avoid compensatory movements, and stand as still as possible. Following the balance test, biomechanical properties of the *gastrocnemius medialis* (MG) and *rectus femoris* (RF) were measured using a handheld, non-invasive MyotonPRO device (Myoton AS, Tallinn, Estonia). Specifically, we measured these muscles ‘dynamic stiffness and dampening (logarithmic decrement) by applying a mechanical impulse and recording the muscle’s response (Garcia-Bernal *et al*., 2021). Five successive measurements were taken for each muscle. Subsequently, the participants were asked to walk wearing their shoes on an instrumented treadmill at 1.11 m s^-1^ (4 km h^-1^). The selected walking speed was similar to the one used in our previous studies (Dewolf *et al*., 2019*b*, 2021*b*, 2021*c*, 2022*b*) and was reported as the comfortable and economical average walking speed based on the net energy consumption (Cavagna & Kaneko, 1977). Before starting the data collection, participants had the opportunity to carry out several tries on the treadmill with oral feedback from the experimenter to familiarize themselves with the task. Data were recorded for 30 seconds after at least 30 seconds of steady-state walking, and on average, 28.1 ± 2.9 (mean ± SD) strides were analysed for each participant.

After completing the walking trial, the participants performed Isometric Maximal Voluntary Torque (IMVT) measurements for the knee and ankle extensors using a custom-made strain gauge following the design of Heglund (1981). The IMVT protocol began with a warm-up, during which participants performed incremental repetitions. Following the warm-up, the participants were instructed to exert their maximum effort for 6 seconds. Three trials were recorded, and the highest torque was analysed. The IMVT data were sampled at 500 Hz. For the knee extensor torque measurements, participants were seated with their hips, knees and ankles positioned at a 90° angle, and a rigid rope was fixed horizontally between the ankle and the gauge fixed on the wall. For ankle extensor torque, the participants were seated in the same position, but with the shank fixed vertically. A rigid rope was fixed vertically between the fifth metatarsal and the strain gauge, fixed on the wall. Participants were instructed to avoid trunk flexion to ensure accurate torque measurements.

The treadmill used for gait and balance analysis (H/P Cosmos-Stellar treadmill, Germany, belt surface: 1.6 x 0.65 m) was equipped with four force transducers (Arsalis®, Belgium). Since the transducers were placed under the body of the treadmill, the force transducers measured the three components of ground reaction force (GRF) exerted by the treadmill belt under the foot (Willems & Gosseye, 2013): F_v_, F_f_, and F_l_, respectively the vertical, fore-aft, and lateral components of the GRF. Data were sampled at a frequency of 1000 Hz.

Electromyographic (EMG) muscle activity from 12 muscles on the right side of the body was captured at a frequency of 2 kHz using a Delsys Trigno Wireless System (Boston, MA). The muscles registered were the *erector spinae* (ES) at the L2 level, the *tensor fasciae latae* (TFL), the *gluteus medius* (Gmed), the *vastus medialis* (VM), the *vastus lateralis* (VL), the *rectus femoris* (RF), the *tibialis anterior* (TA), the *semitendinosus* (ST), the long head of *biceps femoris* (BF), the *gastrocnemius lateralis* (GL), the *gastrocnemius medialis* (GM), and the *soleus* (SOL). Prior to electrode placement, the skin was prepared by shaving and cleaning with fine sandpaper, ether, and alcohol following guidelines from SENIAM (the European project on surface EMG-seniam.org). To ensure accurate placement, the muscle bellies were located by palpation, and electrodes were aligned with the main direction of the muscle fibres (Kendall *et al*., 2005). Electrode placement and signal quality were verified through visual inspection of EMG signals while participants contracted each muscle. EMG data and kinetic measurements were synchronized using a Delsys Trigger Module. Bilateral, full-body three-dimensional kinematics were recorded using a Qualisys system (Gothenburg, Sweden) equipped with 14 Oqus 600+ cameras and 4 Miqus M3 cameras placed around the treadmill. Participants were equipped with 20 retro-reflective markers, placed bilaterally on specific anatomical landmarks: neck, shoulders, wrists, posterior superior iliac spines, anterior superior iliac spines, greater trochanters, knees, malleoli, elbows, and the fifth metatarsal bones, according to the Qualisys sports marker set disposition.

### Data analysis

*Division of the stride:* The foot contact (FC) and toe-off (TO) events were estimated from the displacement of the centre of pressure on the belt (Meurisse et al., 2016). A stride was delimited by two successive right FCs. A step was defined as the interval between one leg’s FC and the contralateral leg’s FC. Stance phases were measured as the time between FC and TO of the same leg.

*Posture:* The lateral (CoP_x_) and anteroposterior (CoP_y_) displacement of the centre of pressure (CoP) were analysed during 30 seconds standing with closed eyes. A 4th-order low-pass Butterworth filter was applied with a 20 Hz cut-off frequency (Dewolf *et al*., 2021*a*).

Specifically, the amplitudes of COP*_x_* and COP*_y_* were determined as two standard deviations (±1 s.d.) of the time series.

*Walking kinetics:* Data were recorded at a sampling rate of 1000 Hz. The fore-aft and vertical velocity of the CoM were determined from the fore-aft and vertical components of the GRF using the procedure described in detail in Dewolf et al. (2016). In short, the fore-aft acceleration of the CoM was calculated as a_f_= F_f_/m, where m is the subject’s body mass. The vertical acceleration of the CoM was calculated as a_v_= (F_v_-m *g*)/m, where *g* is the acceleration of gravity. The vertical (V_v_) and the forward velocity (V_f_) of the CoM were calculated by time- integration of a_v_ and a_f_, respectively, plus an integration constant, which was computed so that the average velocity over a stride was equal to zero. The vertical and forward displacements of the CoM (S_v_ and S_f_, respectively) were then computed by time-integration of V_v_ and V_f_.

The energy of the CoM (E_com_) was computed as the algebraic sum at each instant of its gravitational potential energy (E_p_= m *g* S_v_) and its kinetic energy (E_k_= ½ m (V ^2^ +V_v_ ^2^)). The work done to move the CoM relative to the surroundings (W_com_) was then computed as the sum of the positive increments of E_com_ (Cavagna et al., 1976). The energy transduction between E_p_ and E_k_ was estimated from the relative amount of energy saved over a step (%*recovery*):

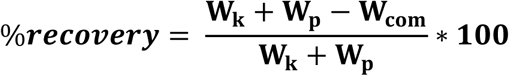

The external power P_com_ was computed as P_com_ = *d*E_com_/*d*t. During a walking stride, the P_com_ time-curve typically shows one major negative peak (W^-^) and two major positive peaks (W^+^ and W^+^), known as collision and propulsion, respectively (Dewolf *et al*., 2019*a*). The maximum values of these peaks were measured for all trials.

*Kinematics:* Kinematic data were recorded at a sampling rate of 240 Hz and later oversampled during post-processing to 1 kHz, to match the kinetic sampling frequency. The elevation angle of each segment (trunk, thigh, shank, and foot), defined as their orientation relative to the vertical (Borghese *et al*., 1996; Dewolf *et al*., 2018), was measured in the sagittal plane. From these angles, the hip, knee, and ankle joint angles were calculated. The hip, knee and ankle joint angle during standing were subtracted from the joint measured during walking tasks (0° corresponds to the standing position). Each stride was interpolated over 400 points, time-normalizing the data across all trials for each subject.

*EMG:* The 12 raw EMG signals were recorded at a sampling rate of 2048 Hz and resampled at 1000 Hz to match the sampling frequency of the kinetic data. The signals were high-pass filtered at 30 Hz, then rectified, and subsequently, low-pass filtered using a zero-lag 4th-order Butterworth filter at 10 Hz. Additionally, a semi-automatic custom procedure was used to detect artefacts by comparing the EMG envelope of each gait cycle with the average envelope across all strides. The time scale was normalized by interpolating each gait cycle to 400 points. For each condition, the full width at half maximum (FWHM) was determined as the duration during which the EMG activity exceeded half of its maximum value (Martino *et al*., 2014; Santuz *et al*., 2020). The centre of activity (CoA) of each EMG signal was also calculated based on the vector’s angle that points to the centre of mass of the circular distribution (Dewolf *et al*., 2022*a*). The CoA was preferred, as identifying a clear peak of activity was impractical for most muscles (Martino *et al*., 2014).

The EMG activity was then mapped onto the rostro-caudal positions of the motor neuron (MN) pools in the human spinal cord, ranging from segments L2 to S2, following Kendall’s myotomal charts (Kendall *et al*., 2005), as used by (Ivanenko *et al*., 2006). The fractional activity value was multiplied by the estimated number of MNs at each segment (MNj), based on the work of Tomlinson & Irving, (1977), to account for the different sizes of MN pools at each level. To compute the total motor output for each condition, the motor output patterns across the gait cycle were summed across the lumbar, sacral and all spinal segments. The mean activation of the lumbar (L2 to L5) and sacral (S1 to S2) segments was computed by averaging the motor output patterns for each region. Finally, the FWHM and the CoA were calculated for both lumbar and sacral segments. The co-activation index (CI) was assessed between the lumbar and sacral segments using the following formula (Rudolph *et al*., 2000; Mari *et al*., 2014):

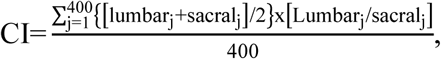

where sacral and lumbar represent the mean activation of the segments during the stride. The CI was then averaged over the entire gait cycle (*j* = 1:400), providing an estimate of the co-activation over the entire cycle. High CI represent a high level of activation of both spinal segments across a large time interval, whereas low CI indicate either a low-level activation of both segments or a high-level activation of one segment along with low-level activation of the other one.

*IMVT:* A custom-made MATLAB script was utilized to accurately determine the peak force generated during the 6-second maximal voluntary contraction (MVC) trial. The script processed the raw data, applying a low-pass filter at 20 Hz. After filtering, the peak torque was identified as the maximum force maintained for at least half a second. This approach avoids the detection of a short peak of force, providing a precise measurement of the participant’s maximal torque output during the isometric test. The lever arms were measured using anthropometers.

### Statistics

For each dependent variable, the normality of the residuals was visually assessed using QQ plots. For variables that did not meet the normality assumption, a log10 transformation was applied, and normality was confirmed afterwards. A mixed-effects model was performed with age (young *vs.* older adults) and physical activity level (less *vs.* more active) as fixed effects. A post-hoc Fisher LSD multiple comparisons test was used to evaluate differences between age groups and levels of physical levels. The interaction between the fixed effect was reported if it was significant. All statistical analyses were conducted using SPSS 23 (IBM, USA) with an alpha level of 0.05.

## Results

Some small differences in subject characteristics were found between groups, presented in Table 1. Subjects physically more active were slightly younger than less active ones (F_1,55_ = 2144.0, *p*< 0.001; F_1,55_ = 6.0, *p*= 0.017). Older adults were on average smaller than young adults (F_1,55_ = 8.6, *p*= 0.004). Regarding the subject’s weight, no significant differences were observed between groups (*p*> 0.743). However, the BMI was slightly smaller in more active individuals (F_1,55_ = 4.0, *p*= 0.049), while age (F_1,55_ = 0.3, *p*= 0.579) had no effect. More importantly, no effects of age (F_1,55_ = 0.9, *p*= 0.334) or interaction (F_1,55_ = 0.4, *p*= 0.511) were observed on the classification using the estimated MET-min/week.

### Muscular properties

The maximal isometric torques were affected by age (Fig. 2B). Indeed, both the knee and ankle extensor strength were smaller in older adults compared to young participants (knee: F_1,55_ = 27.5, *p*< 0.001; ankle: F_1,55_ = 26.9, *p*< 0.001). No difference was observed in knee extensor torque as a function of physical activity level (F_1,55_ = 2.7, *p* = 0.107). However, an effect of level of activity was observed on plantar flexor torque (F_1,55_ = 6.2, *p*= 0.020), with the more active participants being stronger.

**Figure 2.**
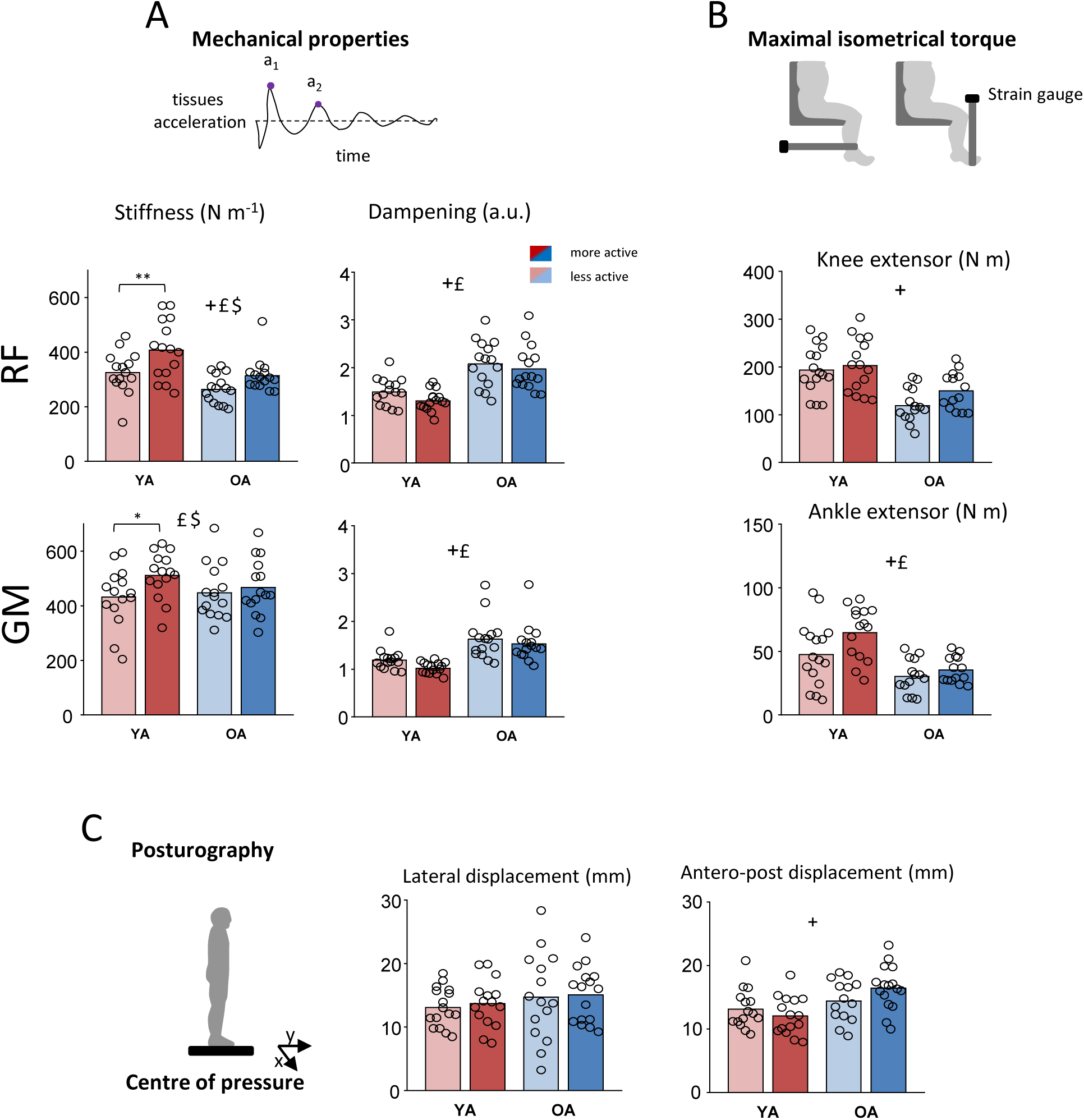
Mechanical properties, maximal isometric torque, and posturography. **A.** Mechanical Properties: The stiffness (N/m) and damping (a.u.) of the *rectus femoris* (RF) and *gastrocnemius medialis* (GM) muscles were measured using the MyotonPRO device. The upper schematic illustrates the tissue acceleration following the brief impulse applied by the MyotonPRO, which records the subsequent oscillatory response of the tissue. The peak a₁ represents the initial acceleration peak and a₂ represents the decaying oscillations as the tissue returns to its resting state. **B.** Maximal Isometric Torque: the schematics illustrate the participant’s position and the orientation of the strain gauge fixation, which varies depending on the muscles being assessed. The bar plots represent the maximal isometric torque of knee extensors and ankle extensors for each group **C.** Posturography: The CoP displacement was measured in both lateral (X) and anteroposterior (Y) directions. The bar plots indicate the average lateral and anteroposterior displacement in each group of participants. In each subplot, the bars represent the group mean, whereas the open circles indicate individual data. A significant effect of age, physical activity level, and their interaction are indicated with the symbols +, £, and $, respectively. The asterisks indicate post hoc comparisons between groups (*p < 0.05, **p < 0.01, ***p < 0.001, ****p < 0.0001).

When comparing the mechanical properties of knee and ankle extensors, the stiffness of both *rectus femoris* and *gastrocnemius medialis* decreased with age (RF: F_1,326_ = 101.8, p < 0.001; GM: F_1,326_ = 5.5, *p* = 0.019). Instead, the tissue dampening increased for both muscles with age (RF: F_1,326_ = 199.0, *p*< 0.001; GM: F_1,326_ = 169.9, *p*< 0.001). The physical activity level significantly influenced both stiffness (RF: F_1,326_ = 76.4, *p*< 0.001; GM: F_1,326_ = 21.7, *p*< 0.001) and tissue dampening (RF: F_1,326_ = 6.8, *p*= 0.009; GM: F_1,326_ = 10.5, *p*= 0.001;). The active participants exhibited higher muscle stiffness and lower tissue dampening. Regarding stiffnesses, the effect of physical activity level was more pronounced in young adults (interaction: RF: F_1,326_ = 8.1, *p*= 0.005; GM: F_1,326_ = 4.9, *p*= 0.027;).

### Static balance

The average centre of pressure (CoP) displacement during standing was compared between groups (Fig. 2C). The lateral displacement of the CoP was not influenced by age or physical activity level (Age: F_1,54_ = 2.4, *p*= 0.121; physical activity level: F_1,54_ = 0.1, *p*= 0.710). Regarding the anteroposterior displacement of the CoP, greater displacement was observed in older adults (F_1,54_ = 4.0, *p*= 0.050), but the level of activity did not affect the static balance (F_1,54_ = 0.56, *p*= 0.457).

### Gait mechanics

Figure 3 illustrates a typical trace of the CoM energy and the time course of the instantaneous recovery within one stride in each group. Older adults walked with shorter steps (F_1,55_ = 32.8, *p*< 0.001). Interestingly, the effect of age was greater for less active participants (physical activity level: F_1,55_ = 8.4, *p*= 0.005; interaction: F_1,55_ = 4.4, *p*= 0.040), with the less active older adults displaying significantly shorter step lengths compared to the more active ones. Similarly, the stance phase was smaller in older adults (F_1,55_ = 29.6, *p*< 0.0001), and in less active participants (F_1,55_ = 4.1, *p*= 0.045).

**Figure 3.**
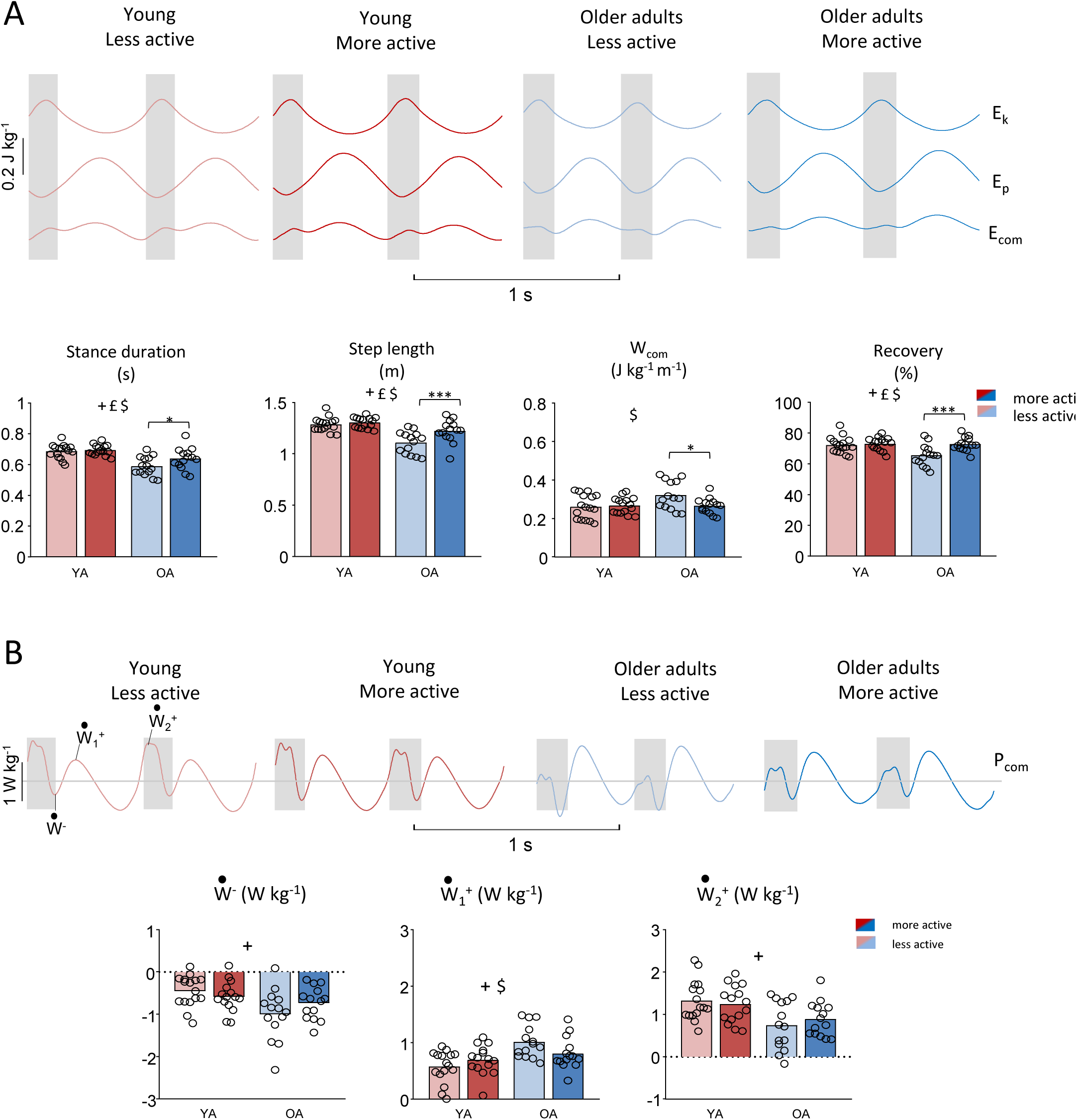
Mechanics of walking. A. Mean traces of kinetic energy (Ek), potential energy (Ep), and total mechanical energy (Ecom) versus time across one representative stride for each group of participants. At the bottom, the bar plots indicate from left to right the average stance duration (s), step length (m), external work to sustain the motion of the CoM (J kg-1 m-1), and the %recovery. B. Mean time curves of the external power of the center of mass (Pcom) in each group of participants. Each trace displays one major negative peak (W⁻, collision) and two major positive peaks, W⁺₁ (weight acceptance) and W⁺₂ (propulsion), during a stride. The bar plots represent the average value of the three peaks of power. In all panels, the grey shaded areas indicate the double-contact phases. The length of the trace along the x-axis represents the mean stride time for each group. Other information as in Fig. 2.

The mechanical work required to sustain the motion of the CoM relative to the surroundings (W_ext_) was not affected by the age (F_1,55_ = 3.8, *p*= 0.056) or physical level of the participants (F_1,55_ = 2.7, *p*= 0.101). However, an interaction between these two factors was found (F_1,55_ = 4.4, p = 0.040), showing that in older adults, the mechanical cost of walking was higher in less active individuals compared to the more active ones (post hoc: *p*= 0.019). Such difference in W_ext_ can partially be explained by a reduction of energy transduction between E_k_ and E_p_. Indeed, the *%recovery* was reduced in older adults (F_1,55_ = 6.3, *p*= 0.014) in less active participants (F_1,55_ = 8.8, *p*= 0.004), with a significant interaction between both fixed factors (F_1,55_ = 5.4, *p*= 0.023). In sum, as for W_ext_ no difference was observed between less and more active young adults (post hoc: p = 0.645), whereas the less active older adults showed a significantly lower *%recovery* than the more active ones (p < 0.001).

The pattern of fluctuation of the COM power during walking (Fig. 3B) typically presents one major peak of negative power (*Ẇ*^−^) and two major peaks of positive power (*Ẇ*^+^& *Ẇ*^+^) (Cavagna *et al*., 1976; Dewolf *et al*., 2019*a*). The three peaks of power were affected by age (*Ẇ*^−^: F_1,55_ = 9.5, *p*= 0.003; *Ẇ*^+^: F_1,55_ = 14.5, *p*< 0.001; *Ẇ*^+^: F_1,55_ = 14.4, *p* < 0.001), with greater *Ẇ*^−^ and *Ẇ*^+^ and smaller *Ẇ*^+^ in older individuals. No significant effect of physical activity level was found (all p> 0.547), but an interaction between age and the level of activity was significant for *Ẇ*^+^ (F_1,55_ = 4.9, *p*= 0.030). The peak of power tended to be greater in older participants less active compared to more active ones, whereas no differences were observed between young adults less or more active (post hoc: *p*= 0.243).

### Gait kinematics

The average lower-limb joint angles (hip, knee and ankle), as well as two orientation angles (trunk and foot), were measured for each group of participants (Figure 4A). In Figure 4B, we have highlighted 3 parameters affected by age: the foot angle at touchdown, the mean trunk angle, and peak knee flexion during stance. Indeed, compared to young adults, the foot at touchdown was less dorsiflexed (F_1,54_ = 4.6, *p*= 0.036), the trunk was more inclined forward (F_1,54_ = 28.2, *p*< 0.0001) and the knee was more flexed during stance (F_1,54_ = 4.7, *p*= 0.033) in older adults. None of these angles was affected by the level of physical activity (all p> 0.201). Nevertheless, an interaction between the effect of age and physical activity level was significant for the foot angle at touchdown (F_1,54_ = 8.5, *p*= 0.005). Indeed, the older adults more active had greater ankle dorsiflexion at touchdown compared to less active ones (post hoc: p = 0.002), whereas no difference was found between groups of younger adults.

**Figure 4.**
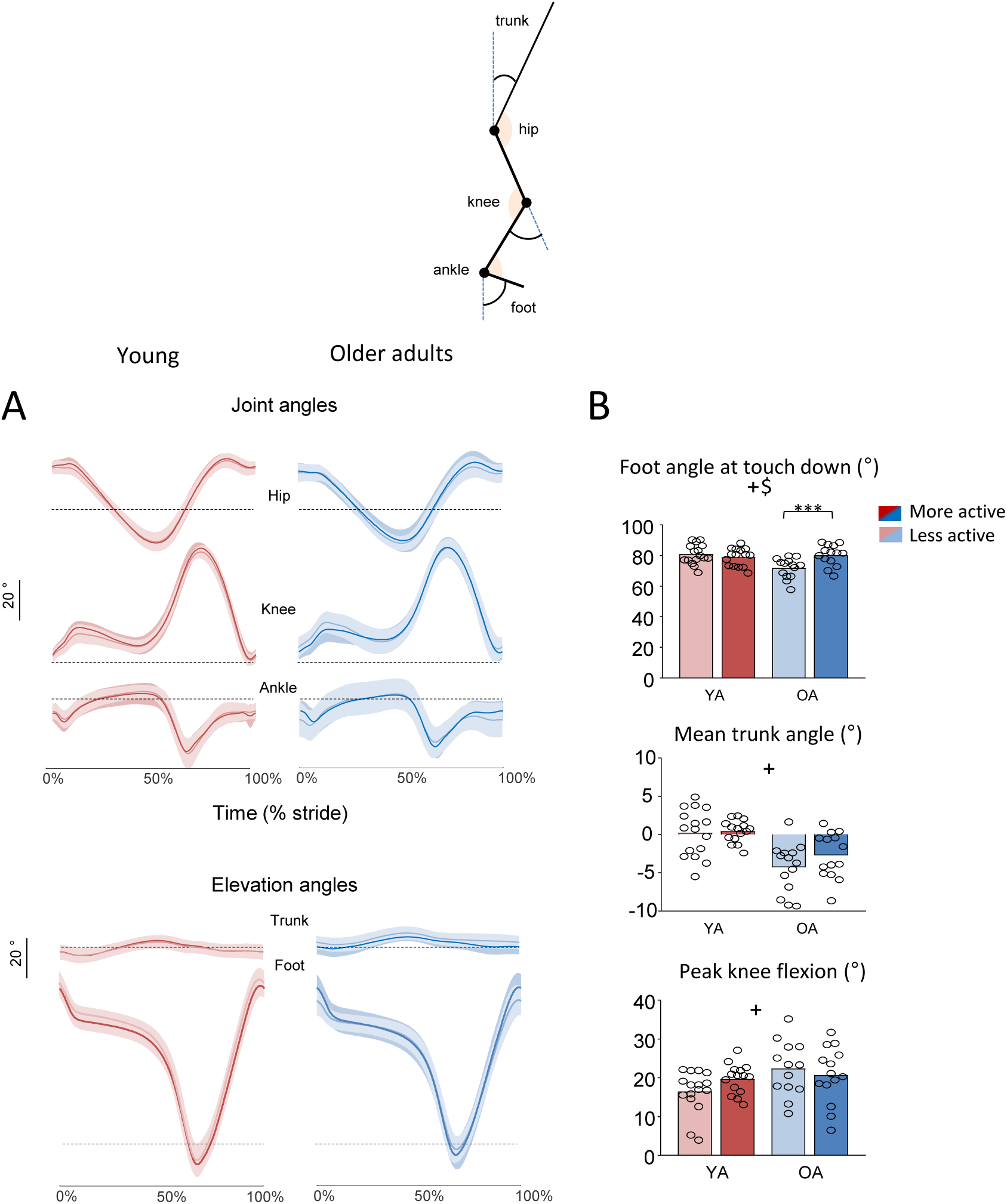
Kinematics of walking. **A.** The stickman illustrates the different joint angles (hip, knee, and ankle) as well as the two orientation angles (trunk and foot) measured in each participant. The mean traces and standard deviations (shaded areas) of each angle over a stride are presented. **B.** The bar plots present the average foot angle at touch down, the mean trunk angle over a stride, and the peak knee flexion during stance in each group. Other indications as in Fig. 2.

### Muscle activity

Figure 5 shows the ensemble averages of rectified EMG envelopes for each group during one walking stride. When comparing the RMS of each EMG, an average greater activation was observed in older adults for all knee extensor and flexor muscles (*vastus medialis, vastus lateralis, rectus femoris, semitendinous, biceps femoris*), and also for *tibialis anterior*, and *gastrocnemius lateralis* (for all F> 6.2 and *p*< 0.016). Also, an effect of physical activity was observed for some muscles (*vastus lateralis*: F_1,53_ = 4.8, *p*= 0.032; *tibialis anterior*: F_1,55_ = 4.0, *p*= 0.049; *gastrocnemius medialis*: F_1,55_ = 6.1, *p*= 0.016), with a higher activation in active individuals. Moreover, for the *erector spinae*, an interaction between age and physical activity level was found (F_1,54_= 5.0, *p*= 0.029), with lower activation in less active older adults compared to their more active peers (post hoc: p = 0.013). For the *tibialis anterior*, post hoc analysis revealed that older adults in the more active group had higher activation than their less active counterparts (*p* = 0.015), while no differences were found in younger adults (p = 0.785).

**Figure 5.**
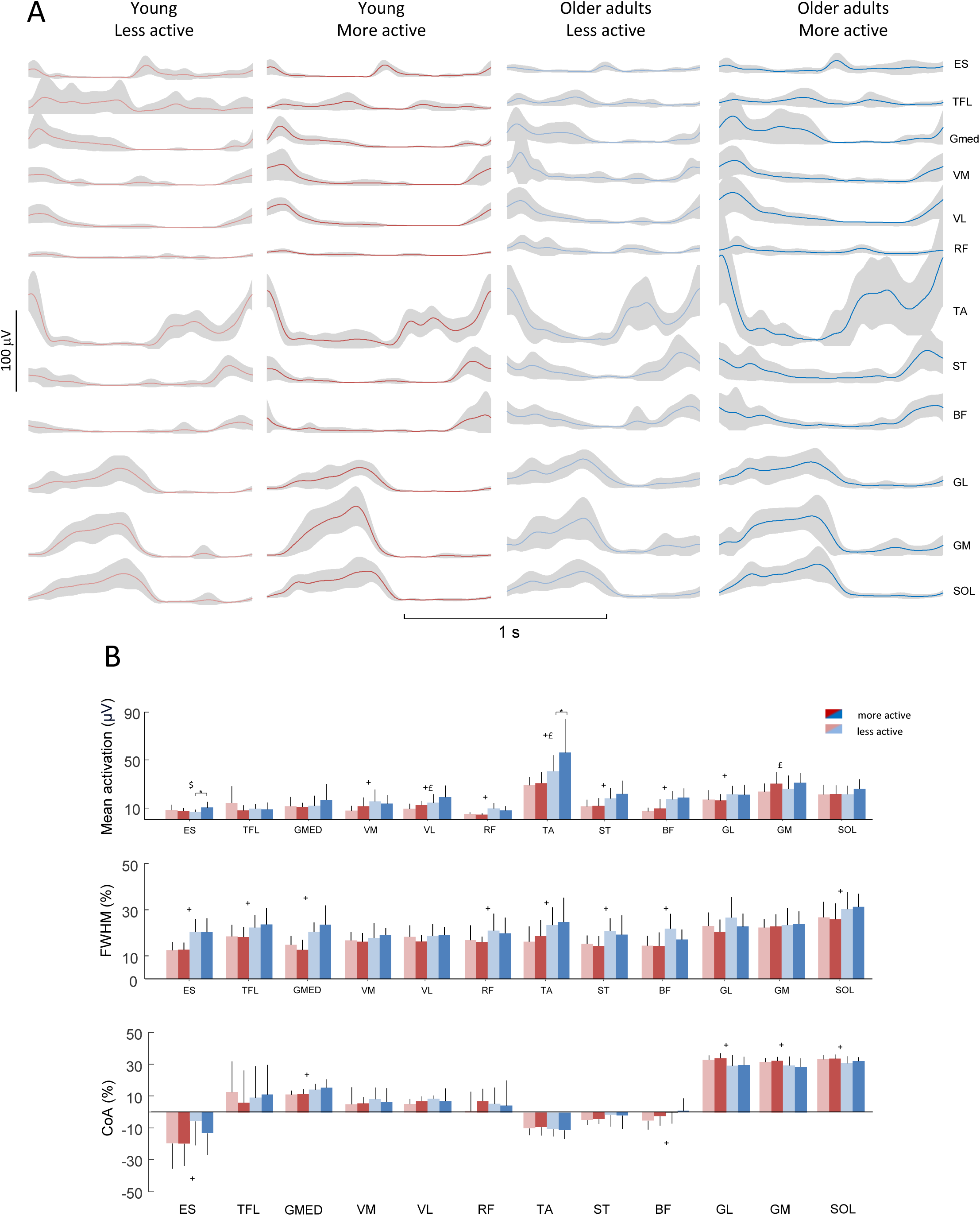
Electromyography (EMG) activation patterns. **A.** Ensemble-averaged EMG curves of the *erector spinae* (ES), *tensor fasciae latae* (TFL), *gluteus medius* (Gmed), *vastus medialis* (VM), *vastus lateralis* (VL), *rectus femoris* (RF), *tibialis anterior* (TA), *semitendinosus* (ST), *biceps femoris* (BF), *gastrocnemius lateralis* (GL), *gastrocnemius medialis* (GM), and *soleus* (SOL). The colored lines represent the ensemble-averaged EMG traces for each group, while the shaded grey areas indicate the standard deviation. **B.** The bar plots represent, from top to bottom, the root mean square (in μV), the full-width half maximum (FWHM, in % of stride), and the center of activation (CoA, in % of stride) for each muscle and each group. The bars correspond to the mean values, with vertical lines above each bar representing one standard deviation. Other indications as in Fig. 2.

Regarding the duration of EMG activation, measured based on the FWHM (Fig. 5B), longer bursts were observed in older adults for a great majority of muscles (ES, TFL, GMED, RF, TA, ST, BF, SOL; for all F> 6.2 and p<0.015). Instead, no effect of physical activity level (or interaction) was found (for all F< 3.3 and p>0.071). The timing of activation was also affected by age. Indeed, the ES, GMED and BF were activated later in the stride in older adults (ES: F_1,54_ = 7.1, *p*= 0.010; GMED: F_1,54_ = 12.8, *p*= 0.001; BF: F_1,53_ = 6.5, *p*= 0.013), whereas the ankle extensors were activated earlier in stance (GL: F_1,55_ = 10.4, *p*= 0.002; GM: F_1,55_ = 7.7, p= 0.007; SOL: F_1,54_ = 5.8, *p* = 0.019). Again, no effect of physical activity level (or interaction) was found on the timing of activation (for all F< 1 and p>0.319).

### Spinal maps

The Figure 6A presents the EMG signals mapped onto the estimated rostrocaudal location of the MN pools in the spinal cord. The timing of activation of the lumbar and sacral motor pools, estimated using the centre of activity (CoA), was different between young and older participants (lumbar: F_1,55_ = 4.8, *p* = 0.032; sacral: F_1,55_ = 22.0, *p* < 0.001). Indeed, the lumbar CoA occurred later whereas the sacral CoA occurred earlier. As a result, the delta between lumbar and sacral activations was reduced in older adults (F_1,55_ = 23.4, *p* < 0.001). While no effect of physical activity was observed on individual muscle activation patterns (Fig. 6), an interaction was found between age and level of physical activity for the lumbar CoA (F_1,55_ = 15.6, *p*< 0.001), with less active older participants displaying a later activation of the lumbar motor pools than more active elders (post hoc: *p* < 0.001), without difference observed between less and more active young participants (post hoc: *p* = 0.082). In turn, the delta between lumbar and sacral CoA was also smaller for less active older adults than more active ones (interaction: F_1,55_ = 8.8, *p*= 0.003; post hoc: p = 0.003).

**Figure 6.**
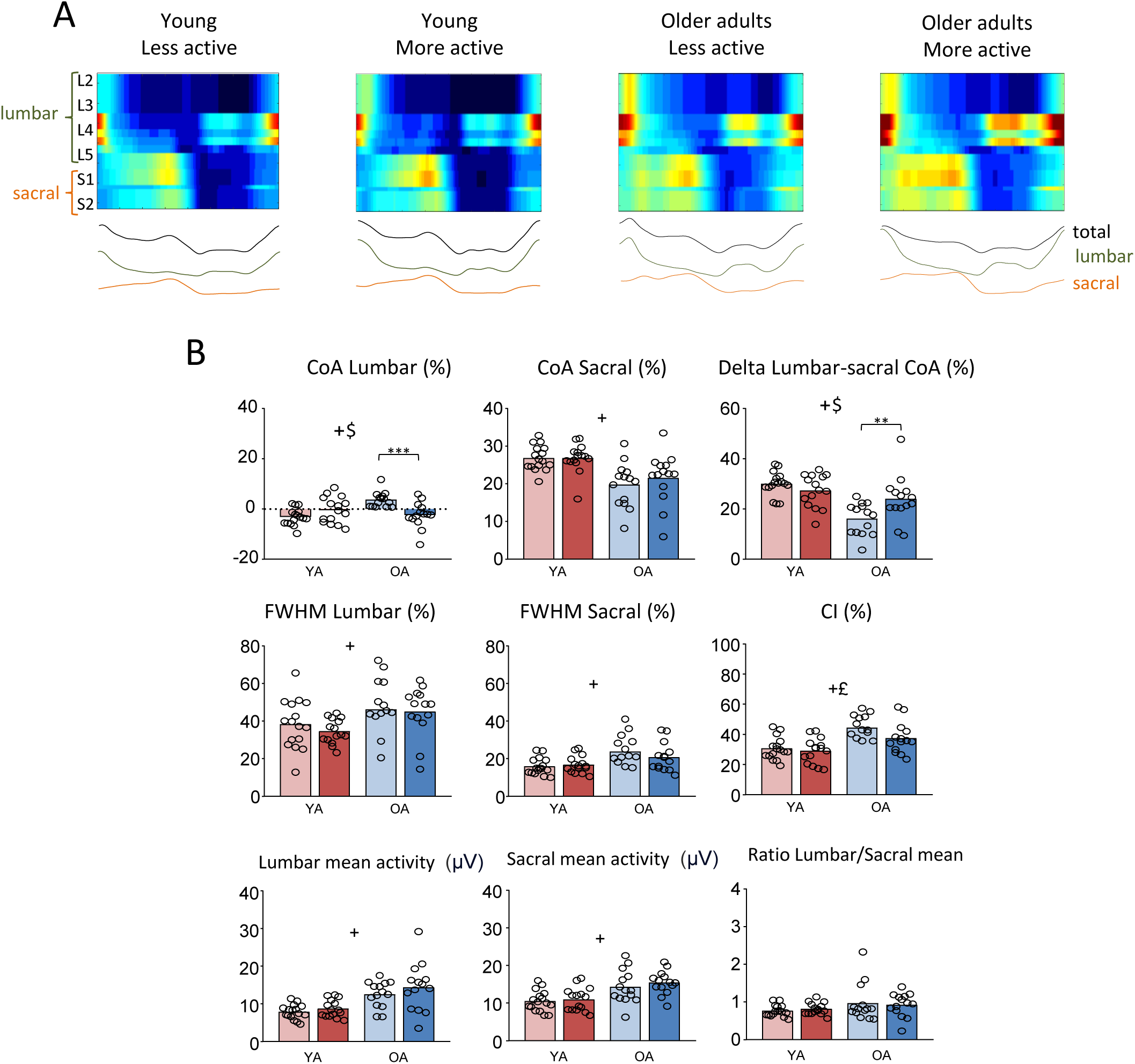
Spinal motor outputs. A. Unilateral spatiotemporal spinal motor outputs computed from ensemble-averaged EMGs for all participants of each group. The total, lumbar, and sacral activation levels are also depicted as traces below the heat maps for each group, relative to the gait cycle. B. Bar plots show the center of activation (CoA, in % of stride) for the lumbar and sacral segments and the delta between lumbar and sacral CoA. Also, the full-width half maximum (FWHM, in % of stride) for both spinal regions, and the coactivation index (CI, %) between lumbar and sacral activations are presented. Finally, the mean activation levels (in μV) of lumbar and sacral outputs, along with the ratio between lumbar and sacral mean activation are displayed. Other indications as in Fig. 2.

The burst durations of the spinal motor output were also affected by age, in both lumbar (F_1,55_ = 12.2, *p*= 0.001) and sacral segments (F_1,55_ = 7.3, *p*= 0.008). Most likely due to longer bursts, the co-activation level (CI) between the lumbar and sacral motor pools was greater in older adults (F_1,55_ = 21.4, *p*< 0.001). While no effect of physical activity level on burst duration was found in both spinal levels (p > 0.05), the co-activation was higher in less active individuals (F_1,55_ = 4.7, *p*= 0.034), with a higher CI in less active older adults than more active ones (post hoc: p = 0.029).

The mean activity of both the lumbar and sacral segments was higher in older adults (F_1,55_ = 26.6, *p*< 0.001; F_1,55_ = 21.8, *p*< 0.001), but the ratio between lumbar and sacral activation was similar in both age groups (F_1,55_ = 3.9, *p*= 0.053). No significant effect was found for the physical activity level in the mean activation of the spinal segments (lumbar: F_1,55_ = 0.7, *p*= 0.616; sacral: F_1,55_ = 0.7, *p*= 0.384) or in the ratio between lumbar and sacral activation (F_1,55_ = 0.002, *p*= 0.966).

## Discussion

In this study, we provide new insights into how a higher level of physical activity can influence the age-related changes observed during walking. Gait in older adults is often characterized by a shorter step length, greater metabolic cost, and distal-to-proximal joint effort redistribution. Our results show that the less active older subjects significantly decrease their step length, increase the power at collision via proximal joints and increase their mechanical work during walking (by about ∼20% relative to young adults; Fig. 3A) relative to more active individuals. While we confirm our hypothesis regarding the impact of aging on muscle mechanical properties and maximal strength, relatively few changes are observed between the most active individuals and the more sedentary ones. However, a modification in spinal motor output was observed in less active individuals, showing an accentuation of the previously documented effect of aging.

### Modification of gait

The metabolic cost of walking is an important variable of daily life that has been studied extensively. In systematic reviews, Das Gupta et al.(2019) and Aboutorabi *et al*. (2016) showed that the average net cost of walking was ∼20% higher in older adults, associated with a loss of independence and an increased risk of morbidity and mortality (Guralnik *et al*., 1995; Cooper *et al*., 2010, 2011; Studenski *et al*., 2011). Previous studies have found that mechanical work does not explain the increase in the metabolic cost of elderly walking since no significant changes have been observed (Mian *et al*., 2006; Ortega & Farley, 2007). Considering only the effect of age, we did not observe any significant change in mechanical work between younger and older individuals (Fig. 3A). However, considering the level of physical activity, we observed a 20% increase in mechanical work in less active older adults. This change can partly be linked to a decrease in the pendular mechanism of walking in less active older adults (Fig. 3A).

In a previous publication, we reported a similar reduction of energy recovery in older adults unable to accelerate the centre of mass forward and upward before double stance because of a lack of trailing leg push-off at the end of stance (Dewolf *et al*., 2021*c*, 2022*b*). Consequently, the centre of mass accelerates after the foot contact of the leading limb (Meurisse *et al*., 2019*a*). Similarly, our results showed that the phase of negative power (*Ẇ*^−^) occurring after collision with the ground, followed by a first phase of positive power (*Ẇ*^+^) were higher in older adults, and specifically in less active ones (Fig. 3B). Both contribute to the vertical redirection of the centre of mass after FC, via greater eccentric energy at the knee extensors of the leading limb (Lay *et al*., 2006; Monaco & Micera, 2012) and were designated as the Collision phase (Dewolf *et al*., 2019*a*). Also, a reduction of the second phase of positive power (*Ẇ*^+^) was reduced in older adults, suggesting a reduction of ankle propulsion in late stance (DeVita & Hortobagyi, 2000; Silder *et al*., 2008). Such distal-to-proximal redistribution with age has garnered considerable scientific attention for nearly 50 years (Winter *et al*., 1990; DeVita & Hortobagyi, 2000; Kerrigan *et al*., 2000; McGibbon, 2003; Franz & Kram, 2013; Franz, 2016; Gueugnon *et al*., 2019; Delabastita *et al*., 2021), most often consider as the hallmark biomechanical ageing features of gait.

While one might expect to observe a change in propulsive force in more active individuals, the age-related reduction of propulsion was similar between groups. Similarly, it has been shown that resistance training consistently improves muscle strength and mitigates sarcopenia but fails to directly translate to improved propulsive power generation in walking (Beijersbergen *et al*., 2013). Instead, the impact of physical activity level was mainly observed during the collision phases (Fig. 3B). Such increase has been associated with deviations from the specific adaptations of the locomotor apparatus to bipedal gait, including the erect posture of the trunk (Núñez-Lisboa *et al*., 2024), straight leg during stance (Cavagna & Kaneko, 1977; Full & Koditschek, 1999), and heel-to-toe rolling pattern of the foot (Usherwood *et al*., 2012; Mesquita *et al*., 2023). Our results showed that less active older adults reduced the heel-to-toe rolling pattern, by decreasing dorsal flexion at foot contact (Fig. 4B). Also, less active individuals tend to increase knee flexion during stance and to increase the average trunk flexion (Fig. 4). All these modifications of kinematic strategy have been related to a greater impact under the leading limb during step-to-step transition (Mesquita *et al*., 2023; Núñez-Lisboa *et al*., 2024), as in older adults. These alterations could stem from age-related declines in neuromuscular function, and the nervous system may adapt by modifying motor strategies to enhance stability and reduce the risk of falls.

### Neuromuscular control of gait

Neural factors are likely to contribute as well to age-related motor impairment (Nùñez-Lisboa *et al*., 2023). For example, changes in control strategy (Peterson & Martin, 2010; Hortobágyi *et al*., 2011; VanSwearingen & Studenski, 2014) have been reported in older adults. Indeed, compared to young adults, older adults exhibited greater activation of lower-limb muscles during walking (Schmitz *et al*., 2009), and higher co-activation of antagonist muscles across the ankle and knee potentially contributing to joint stiffening (Peterson & Martin, 2010; Marques *et al*., 2013). In our results, we observed a greater activation with age in *tibial anterioris* and gastrocnemius lateralis, as well as in knee extensors and flexors (Fig. 5). Also, the timing of muscle activation changed with age, being longer for both ankle flexor (TA) and extensor (SOL) muscles, and knee flexors (ST, BF) and extensor (RF), potentially increasing co-activation.

The greater muscle activation could be related to the decline in muscle quality. Aging muscles weaken (Fig. 2B), which results in an increase in activation and thus in more energy consumed per amount of force produced (Goodpaster et al., 2006; Delmonico et al., 2009). In addition, following the principle of motor unit recruitment, motor unit recruitment progresses from slow-twitch to metabolically more expensive fast-twitch fibres at higher activations (Martin et al., 1992; Bhargava et al., 2004). While the effect of age is in accordance with previous published results, little to no modifications were observed in older adults based on their physical activity level, except for EMG amplitude. Indeed, some muscles (TA, VL, GM and ES) were more active in less active individuals.

Interestingly, insights may be gained by looking into the recruitment and organization of motoneurons during locomotion. The spinal maps, assessed by mapping muscle activity onto the rostrocaudal location of motoneurons, have already revealed specific characteristics of normal aging (Monaco *et al*., 2010; Dewolf *et al*., 2021*b*, 2021*c*; Nùñez-Lisboa *et al*., 2023; Avaltroni *et al*., 2024). As compared to young adults, the motoneurons activation is significantly wider in older adults and the delta between lumbar and sacral activation is reduced. Indeed, the lumbar and sacral activity centres are closer together (Fig. 6), inducing a higher co-activation. Such modification has been related to the lack of late push-off prior to toe-off from the trailing leg (Dewolf *et al*., 2021*c*; Nùñez-Lisboa *et al*., 2023; Avaltroni *et al*., 2024). As for kinematics, it’s interesting to note that we observed a deviation from the two features of motoneuron activation that have been associated with the most energy-efficient bipedal gait: 1) distinct bursts of activity in the lumbar and sacral segments and 2) short burst durations relative to gait cycle duration (Avaltroni *et al*., 2024).

While many age-related changes in gait patterns have been related to physiological deficits in peripheral system function (Horak, 2006), our results suggest that changes in the way the central nervous system coordinates the movements also contribute. In addition, our results show that age-related changes are amplified in older individuals with lower levels of physical activity. For example, the CoA of lumbar motor pools occurred after touch down in less active older adults, as compared to the other groups for whom the activity on average occurred before touch down. It potentially reflects a lack of anticipation before the collision with the ground, as reported earlier in older adults with a higher risk of falls (Meurisse *et al*., 2019*b*). Also, greater co-activation is observed between the activity of the two motoneuron pools in older adults less active, compared to active ones (Fig. 6B). This co-activation reflects the widening of spinal motor output, which has been proposed as a conservative strategy aimed at maintaining gait stability (Martino *et al*., 2015; Santuz *et al*., 2020). This seems to support our hypothesis that engaging in physical activity helps slow down the degradation of the neuromuscular system. Indeed, it’s plausible that regular physical activity may mediate the age-related changes in the central nervous system, and in turn affects movement control strategy (Wang *et al*., 2019; Santuz *et al*., 2020).

### Muscle activity-induced adaptations during aging

The modification of control strategy is also coupled with other intrinsic factors, such as weaker muscles and softer muscle-tendon units in older adults (Monaco & Micera, 2012). Indeed, both knee and ankle extensor maximal isometric torque was reduced with age (Fig. 2B), and individuals with less physical activity displayed reduced ankle extensor maximal forces. Reduced physical activity typically results in skeletal muscle atrophy and an associated reduction of maximal voluntary force (Venturelli *et al*., 2015). Numerous alterations in the physiological properties of the locomotor apparatus have been observed with aging (Pardes *et al*., 2017). Using a predictive neuromechanical model to investigate the physiological origins of age-related gait changes, Song and Geyer (2018) showed that the loss of muscle strength and muscle contraction speed dominantly contribute to the reduced walking economy (Song & Geyer, 2018). The smaller ankle maximal force of less active individuals may exacerbate age differences and induce a greater redistribution of lower extremity joint moment and power compared with more active ones. Such results could explain why vigorous exercises, such as power training or running, prevents the age-related deterioration of muscle (Beijersbergen *et al*., 2017) and, make everyday activities easier (Beck *et al*., 2016).

In addition to strength, age-related changes in the mechanical and morphological properties of lower-limb muscle-tendon units have been reported (Karamanidis & Arampatzis, 2005). Our results also showed that the muscle-tendon stiffness, measured via myotonmetry, was affected by age: the stiffness of rectus femoris and gastrocnemius lateralis was reduced and the dampening (oscillation attenuation) was increased in older adults (Fig. 2A). It has been suggested that age-related changes in neuromuscular activity (higher coactivation) reflect a strategy of stiffening the limb during stance likely to compensate the reduced muscle-tendon stiffness, to utilize tendon elasticity effectively (Gollhofer *et al*., 1992). Indeed, it is suggested that the muscle fascicle behaviours can be related to the motor strategy of older adults to stiffen joints and stabilize motor output in an effort to compensate for reduced muscle strength and for increased tendon compliance and joint laxity (Hortobágyi & DeVita, 2006). Interestingly, a higher level of physical activity in older adults reduced the differences between young and older adults, suggesting a positive impact of the level of activity (Fig. 2A). In addition to the change in strength, this effect of physical activity may contribute to the modification of motor strategy observed between less and more active older individuals, in particular the reduction of co-activation between lumbar and sacral segments (Fig. 6).

### Fallers vs Non-fallers

When comparing older adults according to fall history (non-fallers versus fallers groups), modification of gait and neuromuscular system between groups was similar to the one observed in the present study between less and more active older adults. Indeed, fallers showed shorter stride length (Mortaza *et al*., 2014), increased cost of walking, higher co-activation of lower-limb muscles (Marques *et al*., 2013) and modified motor control strategy (Allen & Franz, 2018). In addition, muscle stiffness of the *gastrocnemius medialis* was significantly lower in the fallers group (Baş *et al*., 2023) and older adult fallers had 28% lower knee extensor strength (Marques *et al*., 2013). Here, we did observe a more variable static balance between young and older participants (Fig. 2C), but no effect of physical activity level. However, given that fallers are presumably less active due to their mobility limitations and fear of falling, it is plausible that their reduced activity levels directly influence these neuromuscular deficits. These findings strengthen our results by highlighting that physical activity plays a crucial role in mitigating age-related changes in gait and neuromuscular function. This reinforces the idea that maintaining or increasing physical activity in older adults can help preserve gait mechanics, and muscle function, and potentially reduce the risk of falls.

### Limitations

The primary limitation of this study is that the level of physical activity was assessed using a self-reported questionnaire, which is inherently less accurate than objective measures such as VO_2_ max (Tangen *et al*., 2022), daily step count (Kraus *et al*., 2019), or accelerometer-based activity tracking (Metcalf *et al*., 2018). These more precise methods could provide a deeper understanding of the relationship between physical activity and gait parameters. However, we believe that our approach also serves as a strength of the study. Despite using a simple classification of activity levels, we observed significant modifications in gait and neuromuscular function related to physical activity. In addition, the individuals classified as ‘less active’ performed an average of 1473.6 METs min/week, exceeding the World Health Organization recommendations (Bull *et al*., 2020). This reinforces the robustness of our findings and supports the main conclusion of the paper, suggesting that even small increases in physical activity could have meaningful effects on gait. For example, transitioning from a lower activity level to a more active group, with just a few additional hours of sports per week, could result in significant improvements in gait mechanics and mobility for less active older adults.

Another limitation of the study is the use of a treadmill. Walking on a treadmill may potentially reduce gait variability as compared to over-ground gait (Lee & Hidler, 2008). However, treadmill walking has been commonly used to investigate age-related modification of gait (Meurisse *et al*., 2019*b*; Dewolf *et al*., 2021*b*, 2022*b*), and spatiotemporal, kinematic, kinetic, muscle activity, and muscle-tendon outcome measures are largely comparable between motorized treadmill and overground locomotion (Van Hooren *et al*., 2020).

## Conclusion

In conclusion, the recognition that the mechanical cost of walking is not purely age-dependent, but rather a consequence of reduced physical activity and subsequent skeletal muscle changes, underscores the vital importance of maintaining physical activity across the lifespan to mitigate the effects of aging on mobility. Indeed, our findings demonstrate that the level of physical activity significantly influences age-related changes in gait and neuromuscular function. While some of these changes are an inherent part of the aging process (Pearson *et al*., 2002), a substantial portion can be attributed to the decline in physical activity levels with advancing age (Hunter *et al*., 2000). Our results underline the critical role of regular physical activity in preserving gait mechanics and muscle function in aging. Therefore, incorporating structured training programs that enhance physical activity levels could be an effective strategy to improve walking performance, reduce walking costs, and delay mobility impairments in older adults (Valenti *et al*., 2016). Such interventions (Malatesta *et al*., 2010) may ultimately contribute to reducing the risk of falls and promoting greater independence as people age.

## Author contributions

The experiments were conducted within the laboratory of physiology and biomechanics of human locomotion – UCLouvain. All authors (A.D. and M.N.) contributed to the conception of the work, the acquisition, analysis and interpretation of data and to writing of the work. All authors approved the final version of the manuscript and agreed to be accountable for all aspects of the work in ensuring that questions related to the accuracy or integrity of any part of the work are appropriately investigated and resolved.

## Acknowledgements

The authors express their gratitude to Antoine Samain and Nemo Lecoubet for their help during the experiments.

## Data availability

The data that support the findings of this study are available from the corresponding author, AHD, upon reasonable request.

## References

1. Aagaard P, Suetta C, Caserotti P, Magnusson SP & Kjaer M (2010). Role of the nervous system in sarcopenia and muscle atrophy with aging: strength training as a countermeasure. Scand J Med Sci Sports 20, 49–64.

2. Aboutorabi A, Arazpour M, Bahramizadeh M, Hutchins SW & Fadayevatan R (2016). The effect of aging on gait parameters in able-bodied older subjects: a literature review. Aging Clin Exp Res 28, 393–405.

3. Akagi R, Yamashita Y & Ueyasu Y (2015). Age-Related Differences in Muscle Shear Moduli in the Lower Extremity. Ultrasound Med Biol 41, 2906–2912.

4. Akima H, Kano Y, Enomoto Y, Ishizu M, Okada M, Oishi Y, Katsuta S & Kuno S-Y (2001). Muscle function in 164 men and women aged 20-84 yr. Medicine and Science in Sports and Exercise 33, 220–226.

5. Allen JL & Franz JR (2018). The motor repertoire of older adult fallers may constrain their response to balance perturbations. J Neurophysiol 120, 2368–2378.

6. Avaltroni P, Cappellini G, Sylos-Labini F, Ivanenko Y & Lacquaniti F (2024). Spinal maps of motoneuron activity during human locomotion: neuromechanical considerations. Front Physiol; DOI: 10.3389/fphys.2024.1389436.

7. Baş H, Okyar Baş A, Ceylan S, Güner M, Koca M, Hafızoğlu M, Şahiner Z, Öztürk Y, Balcı C, Doğu BB, Cankurtaran M & Halil MG (2023). Lower gastrocnemius muscle stiffness, derived from elastography, is an independent factor for falls in older adults. Aging Clin Exp Res 35, 2979– 2986.

8. Beck ON, Kipp S, Roby JM, Grabowski AM, Kram R & Ortega JD (2016). Older runners retain youthful running economy despite biomechanical differences. Medicine and Science in Sports and Exercise 48, 697–704.

9. Beijersbergen CMI, Granacher U, Gäbler M, Devita P & Hortobágyi T (2017). Power Training-induced Increases in Muscle Activation during Gait in Old Adults. Medicine and Science in Sports and Exercise 49, 2198–2205.

10. Beijersbergen CMI, Granacher U, Vandervoort AA, DeVita P & Hortobágyi T (2013). The biomechanical mechanism of how strength and power training improves walking speed in old adults remains unknown. Ageing Research Reviews 12, 618–627.

11. Borghese NA, Bianchi L & Lacquaniti F (1996). Kinematic determinants of human locomotion. J Physiol 494, 863–879.

12. Boyer KA et al. (2023). Age-related changes in gait biomechanics and their impact on the metabolic cost of walking: Report from a National Institute on Aging workshop. Experimental Gerontology 173, 112102.

13. Boyer KA, Andriacchi TP & Beaupre GS (2012). The role of physical activity in changes in walking mechanics with age. Gait and Posture 36, 149–153.

14. Brach JS, Van Swearingen JM, Perera S, Wert DM & Studenski S (2013). Motor learning versus standard walking exercise in older adults with subclinical gait dysfunction: A randomized clinical trial. Journal of the American Geriatrics Society 61, 1879–1886.

15. Bull FC et al. (2020). World Health Organization 2020 guidelines on physical activity and sedentary behaviour. Br J Sports Med 54, 1451–1462.

16. Cavagna GA & Kaneko M (1977). Mechanical work and efficiency in level walking and running. J Physiol 268, 467--81.

17. Cavagna GA, Thys H & Zamboni A (1976). The sources of external work in level walking and running. J Physiol (Lond) 262, 639–657.

18. Çekok FK, Taş S & Aktaş A (2024). Muscle and tendon stiffness of lower extremity in older adults with fall history: Stiffness effect on physical performance and fall risk. Geriatric Nursing 59, 228– 233.

19. Chen N, He X, Feng Y, Ainsworth BE & Liu Y (2021). Effects of resistance training in healthy older people with sarcopenia: a systematic review and meta-analysis of randomized controlled trials. European Review of Aging and Physical Activity 18, 1–19.

20. Cogliati M, Cudicio A, Martinez-Valdes E, Tarperi C, Schena F, Orizio C & Negro F (2020). Half marathon induces changes in central control and peripheral properties of individual motor units in master athletes. Journal of Electromyography and Kinesiology; DOI: 10.1016/j.jelekin.2020.102472.

21. Cooper R, Kuh D, Cooper C, Gale CR, Lawlor DA, Matthews F, Hardy R, & the FALCon and HALCyon Study Teams (2011). Objective measures of physical capability and subsequent health: a systematic review. Age and Ageing 40, 14–23.

22. Cooper R, Kuh D, Hardy R & Group MR (2010). Objectively measured physical capability levels and mortality: systematic review and meta-analysis. BMJ 341, c4467.

23. da Cunha J, Maselli LMF, Stern ACB, Spada C & Bydlowski SP (2015). Impact of antiretroviral therapy on lipid metabolism of human immunodeficiency virus-infected patients: Old and new drugs. World J Virol 4, 56–77.

24. Das Gupta S, Bobbert MF & Kistemaker DA (2019). The Metabolic Cost of Walking in healthy young and older adults – A Systematic Review and Meta Analysis. Sci Rep 9, 9956.

25. Delabastita T, Hollville E, Catteau A, Cortvriendt P, De Groote F & Vanwanseele B (2021). Distal-to-proximal joint mechanics redistribution is a main contributor to reduced walking economy in older adults. Scandinavian Journal of Medicine and Science in Sports; DOI: 10.1111/sms.13929.

26. DeVita P & Hortobagyi T (2000). Age causes a redistribution of joint torques and powers during gait. Journal of Applied Physiology 88, 1804–1811.

27. Dewolf A, La Scaleia V, Fabiano A, Sylos-Labini F, Mondi V, Picone S, Di Paolo A, Paolillo P, Ivanenko Y & Lacquaniti F (2022a). Left–Right Locomotor Coordination in Human Neonates. The Journal of Neuroscience 42, 6566.

28. Dewolf AH, Ivanenko Y, Zelik KE, Lacquaniti F & Willems PA (2018). Kinematic patterns while walking on a slope at different speeds. J Appl Physiol (1985) **125**, 642–653.

29. Dewolf AH, Ivanenko YP, Mesquita RM & Willems PA (2021a). Postural control in the elephant. Journal of Experimental Biology 224, jeb243648.

30. Dewolf AH, Ivanenko YP, Zelik KE, Lacquaniti F & Willems PA (2019a). Differential activation of lumbar and sacral motor pools during walking at different speeds and slopes. J Neurophysiol 122, 872–887.

31. Dewolf AH, Meurisse GM, Ivanenko Y, Lacquaniti F, Bastien GJ & Schepens B (2022b). Relation between Step-To-Step Transition Strategies and Walking Pattern in Older Adults. Applied Sciences 12, 5055.

32. Dewolf AH, Meurisse GM, Schepens B & Willems PA (2019b). Effect of walking speed on the intersegmental coordination of lower-limb segments in elderly adults. Gait & Posture 70, 156–161.

33. Dewolf AH, Peñailillo LE & Willems PA (2016). The rebound of the body during uphill and downhill running at different speeds. Journal of Experimental Biology 219, 2276–2288.

34. Dewolf AH, Sylos-Labini F, Cappellini G, Ivanenko Y & Lacquaniti F (2021b). Age-related changes in the neuromuscular control of forward and backward locomotion. PLOS ONE 16, e0246372.

35. Dewolf AH, Sylos-Labini F, Cappellini G, Zhvansky D, Willems PA, Ivanenko Y & Lacquaniti F (2021c). Neuromuscular Age-Related Adjustment of Gait When Moving Upwards and Downwards. Front Hum Neurosci 15, 749366.

36. Dominici N, Ivanenko YP, Cappellini G, d’Avella A, Mondì V, Cicchese M, Fabiano A, Silei T, Di Paolo A, Giannini C, Poppele RE & Lacquaniti F (2011). Locomotor primitives in newborn babies and their development. *Science* **334**, 997–999.

37. Faul F, Erdfelder E, Lang A-G & Buchner A (2007). G*Power 3: A flexible statistical power analysis program for the social, behavioral, and biomedical sciences. Behavior Research Methods 39, 175–191.

38. Franz JR (2016). The Age-Associated Reduction in Propulsive Power Generation in Walking. Exerc Sport Sci Rev 44, 129–136.

39. Franz JR & Kram R (2013). Advanced age affects the individual leg mechanics of level, uphill, and downhill walking. J Biomech 46, 535–540.

40. Full RJ & Koditschek DE (1999). Templates and anchors: neuromechanical hypotheses of legged locomotion on land. J Exp Biol 202, 3325–3332.

41. Garcia-Bernal M-I, Heredia-Rizo AM, Gonzalez-Garcia P, Cortés-Vega M-D & Casuso-Holgado MJ (2021). Validity and reliability of myotonometry for assessing muscle viscoelastic properties in patients with stroke: a systematic review and meta-analysis. Sci Rep 11, 5062.

42. Gollhofer A, Strojnik V, Rapp W & Schweizer L (1992). Behaviour of triceps surae muscle-tendon complex in different jump conditions. European Journal of Applied Physiology and Occupational Physiology 64, 283–291.

43. Graça AL, Gomez-Florit M, Gomes ME & Docheva D (2023). Tendon Aging. Subcell Biochem 103, 121– 147.

44. Gueugnon M, Stapley PJ, Gouteron A, Lecland C, Morisset C, Casillas J-M, Ornetti P & Laroche D (2019). Age-Related Adaptations of Lower Limb Intersegmental Coordination During Walking. Front Bioeng Biotechnol 7, 173.

45. Guralnik JM, Ferrucci L, Simonsick EM, Salive ME & Wallace RB (1995). Lower-extremity function in persons over the age of 70 years as a predictor of subsequent disability. N Engl J Med 332, 556–561.

46. el Hadouchi M, Kiers H, de Vries R, Veenhof C & van Dieën J (2022). Effectiveness of power training compared to strength training in older adults: a systematic review and meta-analysis. European Review of Aging and Physical Activity 19, 1–15.

47. Harridge SD, Kryger A & Stensgaard A (1999). Knee extensor strength, activation, and size in very elderly people following strength training. Muscle Nerve 22, 831–839.

48. Hassan AS, Fajardo ME, Cummings M, McPherson LM, Negro F, Dewald JPA, Heckman CJ & Pearcey GEP (2021). Estimates of persistent inward currents are reduced in upper limb motor units of older adults. Journal of Physiology 599, 4865–4882.

49. Heglund NC (1981). A simple design for a force-plate measure ground reaction forces. J Exp Biol 93, 333–338.

50. Heppe H, Kohler A, Fleddermann M-T & Zentgraf K (2016). The relationship between expertise in sports, visuospatial, and basic cognitive skills. Frontiers in Psychology; DOI: 10.3389/fpsyg.2016.00904.

51. Hepple RT & Rice CL (2016). Innervation and neuromuscular control in ageing skeletal muscle. Journal of Physiology 594, 1965–1978.

52. Horak FB (2006). Postural orientation and equilibrium: what do we need to know about neural control of balance to prevent falls? Age Ageing 35 Suppl 2, ii7–ii11.

53. Hortobágyi T, A F, S S, P R & P D (2011). Association between muscle activation and metabolic cost of walking in young and old adults. *The journals of gerontology Series A*, Biological sciences and medical sciences; DOI: 10.1093/gerona/glr008.

54. Hortobágyi T & DeVita P (2006). Mechanisms responsible for the age-associated increase in coactivation of antagonist muscles. Exercise and Sport Sciences Reviews 34, 29–35.

55. Hunter GR, Wetzstein CJ, Fields DA, Brown A & Bamman MM (2000). Resistance training increases total energy expenditure and free-living physical activity in older adults. Journal of Applied Physiology 89, 977–984.

56. Hunter SK, Pereira HM & Keenan KG (2016). The aging neuromuscular system and motor performance. J Appl Physiol (1985) **121**, 982–995.

57. Hyatt RH, Whitelaw MN, Bhat A, Scott S & Maxwell JD (1990). Association of muscle strength with functional status of elderly people. Age and Ageing 19, 330–336.

58. Ivanenko YP, Dominici N, Cappellini G, Paolo AD, Giannini C, Poppele RE & Lacquaniti F (2013). Changes in the Spinal Segmental Motor Output for Stepping during Development from Infant to Adult. J Neurosci 33, 3025–3036.

59. Ivanenko YP, Poppele RE & Lacquaniti F (2006). Spinal cord maps of spatiotemporal alpha-motoneuron activation in humans walking at different speeds. J Neurophysiol 95, 602–618.

60. Karamanidis K & Arampatzis A (2005). Mechanical and morphological properties of different muscle-tendon units in the lower extremity and running mechanics: effect of aging and physical activity. J Exp Biol 208, 3907–3923.

61. Kendall F, McCreary E, Provance P, Rodgers M & Romani W (2005). *Muscles. Testing and Function with Posture and Pain*. Lippincott Williams and Wilkins, Baltimore.

62. Kerrigan DC, Riley PO, Rogan S & Burke DT (2000). Compensatory advantages of toe walking. Arch Phys Med Rehabil 81, 38–44.

63. Kraemer WJ, Adams K, Cafarelli E, Dudley GA, Dooly C, Feigenbaum MS, Fleck SJ, Franklin B, Fry AC, Hoffman JR, Newton RU, Potteiger J, Stone MH, Ratamess NA & Triplett-McBride T (2002). Progression models in resistance training for healthy adults. Medicine and Science in Sports and Exercise 34, 364–380.

64. Kraus WE, Janz KF, Powell KE, Campbell WW, Jakicic JM, Troiano RP, Sprow K, Torres A, Piercy KL, & 2018 PHYSICAL ACTIVITY GUIDELINES ADVISORY COMMITTEE* (2019). Daily Step Counts for Measuring Physical Activity Exposure and Its Relation to Health. Med Sci Sports Exerc **51**, 1206–1212.

65. Lay AN, Hass CJ & Gregor RJ (2006). The effects of sloped surfaces on locomotion: A kinematic and kinetic analysis. Journal of Biomechanics 39, 1621–1628.

66. Lee SJ & Hidler J (2008). Biomechanics of overground vs. treadmill walking in healthy individuals. J Appl Physiol (1985) **104**, 747–755.

67. Lexell J (1995). Human aging, muscle mass, and fiber type composition. J Gerontol A Biol Sci Med Sci 50 **Spec No**, 11–16.

68. Lexell J, Henriksson-Larsén K, Winblad B & Sjöström M (1983). Distribution of different fiber types in human skeletal muscles: effects of aging studied in whole muscle cross sections. Muscle Nerve 6, 588–595.

69. Lexell J & Taylor CC (1991). Variability in muscle fibre areas in whole human quadriceps muscle: effects of increasing age. J Anat 174, 239–249.

70. Lim J-Y, Choi SJ, Widrick JJ, Phillips EM & Frontera WR (2019). Passive force and viscoelastic properties of single fibers in human aging muscles. European Journal of Applied Physiology 119, 2339–2348.

71. Malatesta D, Simar D, Saad HB, Préfaut C & Caillaud C (2010). Effect of an overground walking training on gait performance in healthy 65- to 80-year-olds. Experimental Gerontology 45, 427–434.

72. Marcucci L & Reggiani C (2020). Increase of resting muscle stiffness, a less considered component of age-related skeletal muscle impairment. Eur J Transl Myol 30, 8982.

73. Mari S, Serrao M, Casali C, Conte C, Martino G, Ranavolo A, Coppola G, Draicchio F, Padua L, Sandrini G & Pierelli F (2014). Lower limb antagonist muscle co-activation and its relationship with gait parameters in cerebellar ataxia. Cerebellum 13, 226–236.

74. Markov A, Hauser L & Chaabene H (2022). Effects of Concurrent Strength and Endurance Training on Measures of Physical Fitness in Healthy Middle-Aged and Older Adults: A Systematic Review with Meta-Analysis. Sports Medicine 53, 437–455.

75. Marques NR, LaRoche DP, Hallal CZ, Crozara LF, Morcelli MH, Karuka AH, Navega MT & Gonçalves M (2013). Association between energy cost of walking, muscle activation, and biomechanical parameters in older female fallers and non-fallers. Clinical Biomechanics 28, 330–336.

76. Martino G, Ivanenko YP, d’Avella A, Serrao M, Ranavolo A, Draicchio F, Cappellini G, Casali C & Lacquaniti F (2015). Neuromuscular adjustments of gait associated with unstable conditions. J Neurophysiol 114, 2867–2882.

77. Martino G, Ivanenko YP, Serrao M, Ranavolo A, d’Avella A, Draicchio F, Conte C, Casali C & Lacquaniti F (2014). Locomotor patterns in cerebellar ataxia. J Neurophysiol 112, 2810–2821.

78. McGibbon CA (2003). Toward a better understanding of gait changes with age and disablement: neuromuscular adaptation. Exerc Sport Sci Rev 31, 102–108.

79. McNeil CJ, Doherty TJ, Stashuk DW & Rice CL (2005). Motor unit number estimates in the tibialis anterior muscle of young, old, and very old men. Muscle Nerve 31, 461–467.

80. McNeil CJ & Rice CL (2018). Neuromuscular adaptations to healthy aging. Appl Physiol Nutr Metab 43, 1158–1165.

81. McPhee JS, Cameron J, Maden-Wilkinson T, Piasecki M, Yap MH, Jones DA & Degens H (2018). The Contributions of Fiber Atrophy, Fiber Loss, In Situ Specific Force, and Voluntary Activation to Weakness in Sarcopenia. J Gerontol A Biol Sci Med Sci 73, 1287–1294.

82. Mesquita RM, Catavitello G, Willems PA & Dewolf AH (2023). Modification of the locomotor pattern when deviating from the characteristic heel-to-toe rolling pattern during walking. Eur J Appl Physiol; DOI: 10.1007/s00421-023-05169-5.

83. Metcalf KM, Baquero BI, Coronado Garcia ML, Francis SL, Janz KF, Laroche HH & Sewell DK (2018). Calibration of the global physical activity questionnaire to Accelerometry measured physical activity and sedentary behavior. BMC Public Health 18, 412.

84. Meurisse GM, Bastien GJ & Schepens B (2019a). Effect of age and speed on the step-to-step transition phase during walking. Journal of Biomechanics 83, 253–259.

85. Meurisse GM, Bastien GJ & Schepens B (2019b). The step-to-step transition mode: A potential indicator of first-fall risk in elderly adults? PLOS ONE 14, e0220791.

86. Mian OS, Thom JM, Ardigò LP, Narici MV & Minetti AE (2006). Metabolic cost, mechanical work, and efficiency during walking in young and older men. Acta Physiologica 186, 127–139.

87. Monaco V, Ghionzoli A & Micera S (2010). Age-related modifications of muscle synergies and spinal cord activity during locomotion. J Neurophysiol 104, 2092–2102.

88. Monaco V & Micera S (2012). Age-related neuromuscular adaptation does not affect the mechanical efficiency of lower limbs during walking. Gait & Posture 36, 350–355.

89. Mortaza N, Abu Osman NA & Mehdikhani N (2014). Are the spatio-temporal parameters of gait capable of distinguishing a faller from a non-faller elderly? Eur J Phys Rehabil Med 50, 677– 691.

90. Núñez-Lisboa M, Echeverría K, Willems PA, Ivanenko Y, Lacquaniti F & Dewolf AH (2024). Understanding gait alterations: trunk flexion and its effects on walking neuromechanics. Journal of Experimental Biology 227, jeb249307.

91. Nùñez-Lisboa M, Valero-Breton M & Dewolf AH (2023). Unraveling age-related impairment of the neuromuscular system: exploring biomechanical and neurophysiological perspectives. Front Physiol 14, 1194889.

92. Orssatto LBR, Borg DN, Pendrith L, Blazevich AJ, Shield AJ & Trajano GS (2022). Do motoneuron discharge rates slow with aging? A systematic review and meta-analysis. Mechanisms of Ageing and Development; DOI: 10.1016/j.mad.2022.111647.

93. Ortega JD & Farley CT (2007). Individual limb work does not explain the greater metabolic cost of walking in elderly adults. Journal of Applied Physiology 102, 2266–2273.

94. Pardes A, Beach Z, Raja H, Rodriguez A, Freedman B & Soslowsky L (2017). Aging leads to inferior Achilles tendon mechanics and altered ankle function in rodents. J Biomech 60, 30–38.

95. Pearson SJ, Young A, Macaluso A, Devito G, Nimmo MA, Cobbold M & Harridge SDR (2002). Muscle function in elite master weightlifters. Medicine and Science in Sports and Exercise 34, 1199– 1206.

96. Peterson DS & Martin PE (2010). Effects of age and walking speed on coactivation and cost of walking in healthy adults. Gait Posture 31, 355–359.

97. Pour M-AB, Joukar S, Hovanloo F & Najafipour H (2017). Long-term Low-Intensity Endurance Exercise along with Blood-Flow Restriction Improves Muscle Mass and Neuromuscular Junction Compartments in Old Rats. Iran J Med Sci 42, 569–576.

98. Reid KF & Fielding RA (2012). Skeletal Muscle Power: A Critical Determinant of Physical Functioning In Older Adults. Exerc Sport Sci Rev 40, 4–12.

99. Rogers RS & Nishimune H (2017). THE ROLE OF LAMININS IN THE ORGANIZATION AND FUNCTION OF NEUROMUSCULAR JUNCTIONS. Matrix Biol 57–58, 86–105.

100. Rudolph KS, Axe MJ & Snyder-Mackler L (2000). Dynamic stability after ACL injury: who can hop? Knee Surg Sports Traumatol Arthrosc 8, 262–269.

101. Rygiel KA, Picard M & Turnbull DM (2016). The ageing neuromuscular system and sarcopenia: a mitochondrial perspective. Journal of Physiology 594, 4499–4512.

102. Santuz A, Brüll L, Ekizos A, Schroll A, Eckardt N, Kibele A, Schwenk M & Arampatzis A (2020). Neuromotor Dynamics of Human Locomotion in Challenging Settings. iScience 23, 100796.

103. Schmitz A, Silder A, Heiderscheit B, Mahoney J & Thelen DG (2009). Differences in lower-extremity muscular activation during walking between healthy older and young adults. J Electromyogr Kinesiol 19, 1085–1091.

104. Şendur HN, Cindil E, Cerit MN, Kılıç P, Gültekin Iİ & Oktar SÖ (2020). Evaluation of effects of aging on skeletal muscle elasticity using shear wave elastography. Eur J Radiol 128, 109038.

105. Silder A, Heiderscheit B & Thelen DG (2008). Active and passive contributions to joint kinetics during walking in older adults. Journal of Biomechanics 41, 1520–1527.

106. Song S & Geyer H (2018). Predictive neuromechanical simulations indicate why walking performance declines with ageing. Journal of Physiology 596, 1199–1210.

107. Stenroth L, Peltonen J, Cronin NJ, Sipilä S & Finni T (2012). Age-related differences in Achilles tendon properties and triceps surae muscle architecture in vivo. Journal of Applied Physiology 113, 1537–1544.

108. Studenski S, Perera S, Patel K, Rosano C, Faulkner K, Inzitari M, Brach J, Chandler J, Cawthon P, Connor EB, Nevitt M, Visser M, Kritchevsky S, Badinelli S, Harris T, Newman AB, Cauley J, Ferrucci L & Guralnik J (2011). Gait speed and survival in older adults. JAMA 305, 50–58.

109. Tangen EM, Gjestvang C, Stensrud T & Haakstad LAH (2022). Is there an association between total physical activity level and VO2max among fitness club members? A cross-sectional study. BMC Sports Science, Medicine and Rehabilitation 14, 109.

110. Tomlinson BE & Irving D (1977). The numbers of limb motor neurons in the human lumbosacral cord throughout life. Journal of the Neurological Sciences 34, 213–219.

111. Usherwood JR, Channon AJ, Myatt JP, Rankin JW & Hubel TY (2012). The human foot and heel-sole- toe walking strategy: A mechanism enabling an inverted pendular gait with low isometric muscle force? Journal of the Royal Society Interface 9, 2396–2402.

112. Valenti G, Bonomi AG & Westerterp KR (2016). Multicomponent Fitness Training Improves Walking Economy in Older Adults. Medicine and Science in Sports and Exercise 48, 1365–1370.

113. Van Hooren B, Fuller JT, Buckley JD, Miller JR, Sewell K, Rao G, Barton C, Bishop C & Willy RW (2020). Is Motorized Treadmill Running Biomechanically Comparable to Overground Running? A Systematic Review and Meta-Analysis of Cross-Over Studies. Sports Med 50, 785–813.

114. VanSwearingen JM & Studenski SA (2014). Aging, Motor Skill, and the Energy Cost of Walking: Implications for the Prevention and Treatment of Mobility Decline in Older Persons. J Gerontol A Biol Sci Med Sci 69, 1429–1436.

115. Venturelli M, Cè E, Limonta E, Schena F, Caimi B, Carugo S, Veicsteinas A & Esposito F (2015). Effects of endurance, circuit, and relaxing training on cardiovascular risk factors in hypertensive elderly patients. Age; DOI: 10.1007/s11357-015-9835-4.

116. Wang Q, Cui C, Zhang N, Lin W, Chai S, Chow SK-H, Wong RMY, Hu Y, Law SW & Cheung W-H (2024). Effects of physical exercise on neuromuscular junction degeneration during ageing: A systematic review. Journal of Orthopaedic Translation 46, 91–102.

117. Wang Y, Watanabe K & Asaka T (2019). Effect of dance on multi-muscle synergies in older adults: a cross-sectional study. BMC Geriatrics 19, 340.

118. Willems PA & Gosseye TP (2013). Does an instrumented treadmill correctly measure the ground reaction forces? Biol Open 2, 1421–1424.

119. Williamson DL, Godard MP, Porter DA, Costill DL & Trappe SW (2000). Progressive resistance training reduces myosin heavy chain coexpression in single muscle fibers from older men. Journal of Applied Physiology 88, 627–633.

120. Winter DA, Patla AE, Frank JS & Walt SE (1990). Biomechanical walking pattern changes in the fit and healthy elderly. Phys Ther 70, 340–347.

